# In-text citation error rate as a scientometric tool for evaluating accuracy and weighing evidence

**DOI:** 10.1101/2022.06.01.494325

**Authors:** Derek E. Lee

**Affiliations:** Department of Biology, Pennsylvania State University, University Park, Pennsylvania, USA

**Keywords:** cognitive bias, quotation accuracy, citation accuracy, reference accuracy, scientometrics

## Abstract

Scientists are fallable and biased, but accuracy can be assessed through empirical analysis of published work that quantifies in-text citation (or quotation) errors. In scientific conflicts, it can be difficult for outsiders to know whose evidence or interpretation to trust. In-text citation error rate can assist decision- and policy-making bodies, as well as the courts when conflicts reach the judicial branch of government, by quantifying absolute and relative accuracy of scientists presenting scientific evidence. I propose the use of in-text citation error rates as a scientometric tool to quantify the accuracy of an author’s work. In-text citation error rates in excess of an established overall mean (e.g., 11% for minor errors and 7% for major errors in ecology), or differences in in-text citation error rates between opposing groups of scientists could be used to reveal excessive inaccuracies in an author or group. The spotted owl (*Strix occidentalis*) has been at the center of a multi-decadal conflict caused by competition among people over forest resources, with scientific experts representing opposing stakeholders often presenting conflicting evidence. I applied the in-text citation error rate tool to important papers in the spotted owl and forest fire debate and found evidence of greater error rates in works on one side of this debate. In-text citation error rate can be an effective tool for quantifying accuracy among scientists.

## Introduction

During knowledge production, scientists use in-text citations (also called quotations) to connect text statements to previously published material in order to build logical arguments, substantiate assertions, attribute the sources of ideas, or provide important background information for readers (Grafton, 1999). The accuracy of an in-text citation can be defined as the extent to which content of a cited reference supports or is in accordance with the text statement made by the authors (Jergas & Baethge, 2015). Estimates of overall rates of major in-text citation errors (text seriously misrepresenting or bearing no resemblance to the original source) from ecology, marine biology, and medical science were 7.2%, 6.0%, and 11.9%, respectively (Todd et al., 2007; Todd et al., 2010; Jergas & Baethge, 2015). Inaccurate representations of previously published material misleads readers about the findings in the cited source, and interrupts the accurate flow of information to readers (Hosseini et al., 2020). In-text citation errors are widespread (Todd et al., 2007; Todd et al., 2010; Jergas & Baethge, 2015; Smith & Cumberledge, 2020), and can propagate and reinforce such that unfounded claims may become accepted as facts (Greenberg et al., 2009).

Scientists, like all humans, are biased (Keohler, 1993; MacCoun, 1998; Kaptchuk, 2003; Pannucci & Wilkins, 2010), and make mistakes (Todd et al., 2007; Jergas & Baethge, 2015; Smith & Cumberledge, 2020). The accuracy of scientific statements is quantifyable through empirical analysis of in-text citation errors in published scientific texts. One way to classify in-text citation errors is to grade them as major or minor (Todd et al., 2007: Jergas & Baethge, 2015). A major in-text citation error is when the source referred to is not at all in accordance with the claim of the authors (Todd et al., 2007; Jergas & Baethge, 2015). Minor errors are defined as inconsistencies and factual errors not severe enough to be considered contradictory (Todd et al., 2007; Jergas & Baethge, 2015). Minor errors include text that could mislead a reader but do not fundamentally alter the meaning of the source (De Lacey et al., 1985).

In scientific conflicts, it can be difficult for outsiders to understand the science or know whose evidence or interpretation to trust (Krimsky, 2005; Edmond, 2008; Bromme et al., 2015; Sinatra & Hofer, 2016). A growing literature makes the case for evidence-based conservation decision-making (e.g., Pullin & Knight, 2001; Sutherland et al., 2004; Adams & Sandbrook, 2013). However, even when systematic metaanalysis provides strong evidence-based policy guidance (e.g., Lee, 2018; Lee, 2020), there can be substantial push-back by scientists who, for myriad possible reasons, voice opposition to that policy guidance (Jones et al., 2020), creating uncertainty among decision makers, policy makers, judges, and the public.

I propose the use of in-text citation error rates as a scientometric tool to quantify the accuracy of an author’s work. In-text citation error rates in excess of an established overall mean (e.g., 11% for minor errors and 7% for major errors in ecology; Todd et al., 2007), or differences in in-text citation error rates between opposing groups of scientists could be used to reveal excessive inaccuracies in an author or group. As a worked example of this scientometric tool, I evaluated papers from both sides of an ongoing scientific conflict about the effects of mixed-severity forest fire on spotted owls (*Strix occidentalis*) to determine which side is more accurate in their in-text citation practices. Results should help inform and guide appropriate actions for forest management.

## Methods

I evaluated 4 of the most important papers in the ongoing scientific conflict over spotted owls and forest fire, 2 from each side of the issue (Bond et al., 2009; Jones et al., 2016; Jones et al., 2020; Lee, 2020). Forest fire is a natural disturbance event (Marlon et al., 2012) and prominent management issue that can affect habitat of spotted owls, a species that has driven many forest-management decisions over the past few decades (Weatherspoon et al., 1992; USDA, 2018). In summary, Bond and Lee claim/argue/have concluded that wildfire is not a threat to spotted owl population persistence so logging should not be used to alter vegetation in spotted owl habitat, while Jones et al. claim/argue/have concluded wildfire harms spotted owls, so logging spotted owl habitat to reduce fire severity and extent is justified (Jones et al., 2020).

I treated each paper as an independent sample, and compared mean error rates among three groups: 1) Bond and Lee, 2) Jones et al., and 3) mean ecological error rates (11% for minor errors and 7% for major errors) from Todd et al. (2007). I computed in-text citation error (also called quotation error) rate as a ratio assigned to each published article. The denominator was the number of citations examined in the text of the article. The numerator was the number of erroneous in-text citations. I was not concerned with bibliographic errors such as author name spelling or page references.

I randomly selected 20 in-text citations from each article. Every citation was evaluated by 3 independent readers as to whether it was a primary scientific source that presented data and results that directly supported the text statement. A major citation error was when the source referred to is not at all in accordance with the claim of the authors (Todd et al., 2007; Jergas & Baethge, 2015). Minor errors were defined as inconsistencies and factual errors not severe enough to be considered contradictory (Todd et al., 2007; Jergas & Baethge, 2015). Minor errors include text that could mislead a reader, but did not fundamentally alter the meaning of the source (De Lacey et al., 1985). Readers evaluated each citation independently as to whether a minor or major citation error existed, and final evaluation regarding each citation’s accuracy was decided by majority. If a text claim cited to a source where the text claim also appeared, I examined the cited primary material to determine whether and to what degree it supported the claim. Data were collected in a speadsheet for analyses and are available as Supplementary Material (**Table S1**).

## Results

In-text citation error rates of Jones et al. were greater than both mean ecology error rates and Bond & Lee error rates for both minor and major errors (**Fig. 1**). Major error rates of Jones et al. were 3.75x greater than major error rates of Bond & Lee, and 4.3x greater than the mean major error rate of ecology journals. Minor error rates of Jones et al. were 2x greater than error rates of Bond & Lee, and 1.8x greater than mean ecology journal minor error rates. Minor and major in-text citation error rates of Bond & Lee were nearly equivalent to mean ecology error rates.

**Figure 1.**
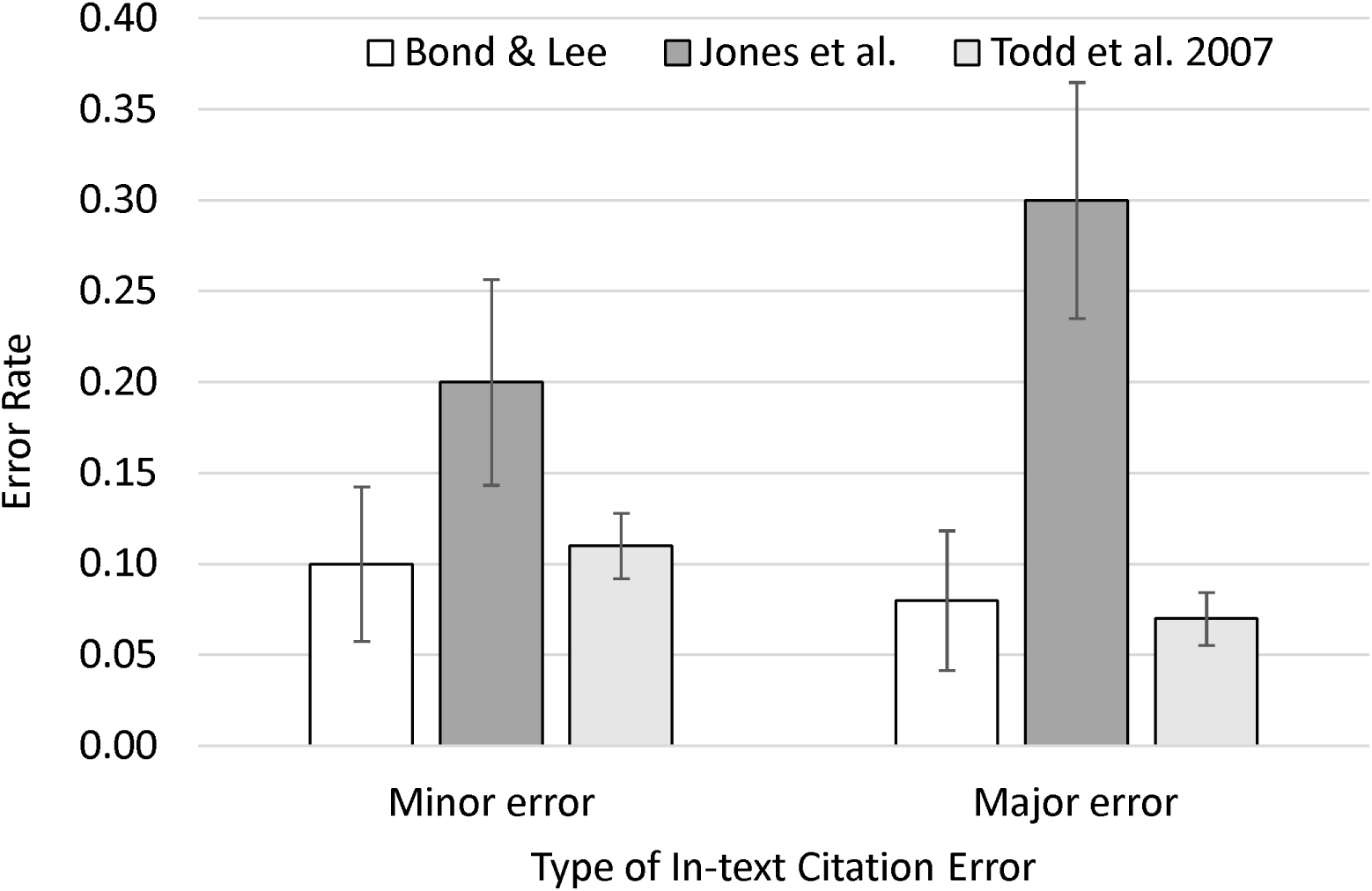
In-text citation error rates, a scientometric quantity used to evaluate accuracy, from two groups of authors on opposing sides of the spotted owl (*Strix occidentalis*) and wildfire debate (group 1 is Bond & Lee, group 2 is Jones et al.), compared with mean ecological in-text citation error rates (Todd et al. 2007). Bond & Lee claim wildfire is not a threat to spotted owls, while Jones et al. claim wildfire is a threat to spotted owls. Jones et al.’s major error rate was 4.3x greater than mean ecology error rates, and 3.7x greater than the major error rate of Bond & Lee. Error bars are standard errors of proportions.

## DISCUSSION

In-text citation error rate can be an effective tool for quantifying absolute and relative accuracy of sources of scientific evidence. In-text citation error rate can illustrate accuracy of authors via comparisons with overall error rates in a field, and relative error rates between groups can quantify relative accuracy of opposing groups of scientists. One strength of this tool is that in an arena of ongoing scientific debate, such as spotted owl responses to forest fire, the antagonists will inevitably be citing the same pool of primary source materials, so deviations from accurate citation practices serve to demonstrate which authors are more accurately interpreting those materials. Such information can be important for decision makers to better understand scientific results to guide appropriate forest management practices. In this case, forest managers should be alerted that citation errors from Jones et al. reveal biases about adverse impacts of forest fires on spotted owls.

Decades of research on cognitive biases has shown that people seek out and evaluate information in ways that favor their preexisting beliefs (Rosenthal, 1976; Kunda, 1990; Nickerson, 1998). Humans prefer information that supports their prexisting beliefs while ignoring or discrediting information incompatible with their beliefs (Kleck & Wheaton, 1967; Koriat et al., 1980). Scientists are not immune to this cognitive trait (Pannuci & Wilkins, 2008), but awareness of biases in science and of effective methods to address them is low among ecology scientists (Zvereva & Kozlov, 2021). Acknowledging the potential for biases of participants in a resource conflict may be useful to decision makers when evaluating the relative strength of opposing claims.

In-text citations are important to the process of knowledge production and should be carefully curated by authors, editors, reviewers, and readers (for more details on the recommendations below see Jergas & Baethge, 2015; Hosseini et al., 2020). In general, authors should verify their statements against original sources, not indirect sources such as reviews. If reviews are used, state it explicitly (as reviewed in Citation). Authors should also place citations immediately after the relevant text phrase, not at the end of a sentence. Editors and reviewers should check at least a sample of citations for accuracy. Readers should be skeptical of citations and alert journals whenever citation errors are discovered. Journals should institute and scientists should conduct post-publication reviews that allow errors to be highlighted or corrected.

## Statements and Declarations

The author did not receive support from any organization for the submitted work. The author declares they have no financial interests.

## Statements and Declarations

The author has no relevant financial or non-financial interests to disclose.

## Data Availability

Data are available as Supplementary Material (Table S1).

